# Connectivity and dynamics in the olfactory bulb

**DOI:** 10.1101/2021.07.19.452784

**Authors:** David E. Chen Kersen, Gaia Tavoni, Vijay Balasubramanian

## Abstract

Dendrodendritic interactions between excitatory mitral cells and inhibitory granule cells in the olfactory bulb create a dense interaction network, reorganizing sensory representations of odors and, consequently, perception. Large-scale computational models are needed for revealing how the collective behavior of this network emerges from its global architecture. We propose an approach where we summarize anatomical information through dendritic geometry and density distributions which we use to calculate the probability of synapse between mitral and granule cells, while capturing activity patterns of each cell type in the neural dynamical systems theory of Izhikevich. In this way, we generate an efficient, anatomically and physiologically realistic large-scale model of the olfactory bulb network. Our model reproduces known connectivity between sister vs. non-sister mitral cells; measured patterns of lateral inhibition; and theta, beta, and gamma oscillations. It in turn predicts testable relations between network structure, lateral inhibition, and odor pattern decorrelation; between the density of granule cell activity and LFP oscillation frequency; how cortical feedback to granule cells affects mitral cell activity; and how cortical feedback to mitral cells is modulated by the network embedding. Additionally, the methodology we describe here provides a tractable tool for other researchers.

**Author summary:** The function of the olfactory bulb (OB) critically depends on connectivity patterns between its excitatory and inhibitory cells. Here, we develop an anatomically grounded algorithm for efficiently determining the probability of synapses between mitral cells and granule cells in the OB. We use this algorithm to generate a large-scale network model of the OB with characteristic connectivity distributions between cell types, as well as between sister mitral cells. We simulate the network using the dynamical systems approach of Izhikevich for describing neurons, and show how network structure affects GC-mediated processes, including LFP oscillation frequency, lateral inhibition, odor decorrelation, and cortical feedback. Our results suggest how alterations to the OB network through processes like neurogenesis, or via injury or disease, can have significant effects on function.

## Introduction

The olfactory bulb (OB), an important waystation along the olfactory pathway, synthesizes odor input with feedback from higher cortical structures via its complex internal circuitry. This synthesis occurs through interactions between two principal components of the bulb, excitatory mitral cells (MCs) and inhibitory granule cells (GCs), which create a network that reshapes odor information as it passes to cortex. Computational studies are necessary for understanding how this odor information is reshaped, since we lack experimental methods for interrogating this network’s structure-function dependency. However, the sheer number of neurons, encompassing tens of thousands of excitatory cells and millions of inhibitory cells [1]; intricate network architecture [2–4]; and complex spiking dynamics [5–9] make detailed biophysical simulation impractical at large scale. Thus, many studies use random connections or simple distance-dependent functions to establish MC-GC connectivity [10–22] and often study smaller networks on the order of hundreds or even tens of neurons [10, 11, 13, 14, 19, 22–24], allowing for highly complex, conductance-based neuronal frameworks [7]. Other approaches use rate-based or population equations [12, 21, 25] thereby facilitating models with larger numbers of neurons. Together, these studies have shed light on important OB phenomena, such as beta and gamma oscillations [11, 13–15, 17, 20] and effects associated with olfactory discrimination and perceptual learning [12, 19, 21, 26–28].

However, OB function depends heavily on connectivity: particular arrangements of GCs around MCs can dramatically affect its output [8, 29–38]. Likewise, the characteristic spiking dynamics of GCs and MCs can change overall network behavior, e.g., affecting the nature of oscillations that may play an important role in olfactory coding and perception [14, 39]. With this in mind, we have leveraged diverse anatomical and physiological data to build on earlier models and craft an algorithm for generating large-scale, realistic networks of MCs and GC that facilitate studies of emergent OB network behavior.

In short, we inferred the average distribution of dendrites for each cell type from data [8, 40, 41] and modeled the results geometrically. By calculating the intersection between the dendritic distributions of a given MC and GC, we could extrapolate the average number of synapses for the cell pair and in turn calculate a probability of synapsing. After constructing spatial distributions of MCs and GCs in the OB, we sampled the synapse probability for each MC-GC pair to build a large-scale network constrained by the anatomy and featuring a realistic ratio of GCs to MCs [1]. We modeled each cell using the dynamical systems theory developed by Izhikevich, thus reproducing realistic cellular spiking patterns [8, 9, 42, 43]. The resulting network was tractable: a network with nearly 20,000 units could be simulated for tens of thousands of steps (several seconds of real time) in a few hours on a conventional laptop; parallelizing on a server will divide simulation time by roughly the number of processors used.

Our model reproduced important empirical features of the OB, including differential connectivity patterns among sister and non-sister MCs, decorrelation over short timescales, as well as theta, beta and gamma oscillations in the local field potential (LFP). The model makes the surprising, and testable, prediction that cortical feedback inhibition of MCs via GCs is a network property largely independent of which GCs are targeted, an observation with consequences for our understanding of how context modulates odor representations [28, 44–53], and for theories of the functional purpose of granule cell neurogenesis [12, 21, 26, 36]. The model also predicts that beta and gamma oscillations, which are implicated in numerous theories of odor coding and decoding [54–57], are network properties intrinsic to the bulb that can be modified, suppressed, or enhanced by the density of granule cell activity [14, 23, 39].

## Results

### Deriving the probability of synapse

To establish the probability that a particular mitral cell (MC) and granule cell (GC) form a synapse, we first determined the average number of synapses between that pair. Unlike most neurons, MCs and GCs of the olfactory bulb (OB) form synapses between dendrites, specifically between MC lateral dendrites, which extend out from the soma along the contour of the bulb, and GC dendritic spines, small ellipsoid protuberances off the GC dendrite [58, 59]. The dendritic trees of these cells have stereotypical shapes, so we approximated the OB as a flat 3-dimensional space, and used simple geometric forms to represent the mean spatial distributions of these trees (Fig. 1A). The lateral dendrites of MCs extend radially, roughly in a disk when viewed from above [9, 40, 41], so we defined the MC dendritic tree as a radially-symmetric distribution on a flat disk, with each disk oriented parallel to the faces of the OB space. *Camera lucida* images from [40] and [41], showed that the density (in *µ*m of lateral dendrite per *µ*m^2^ area) at a given radial distance *r* from the soma was well fit by the function (Fig. 1C):

**Fig 1.**
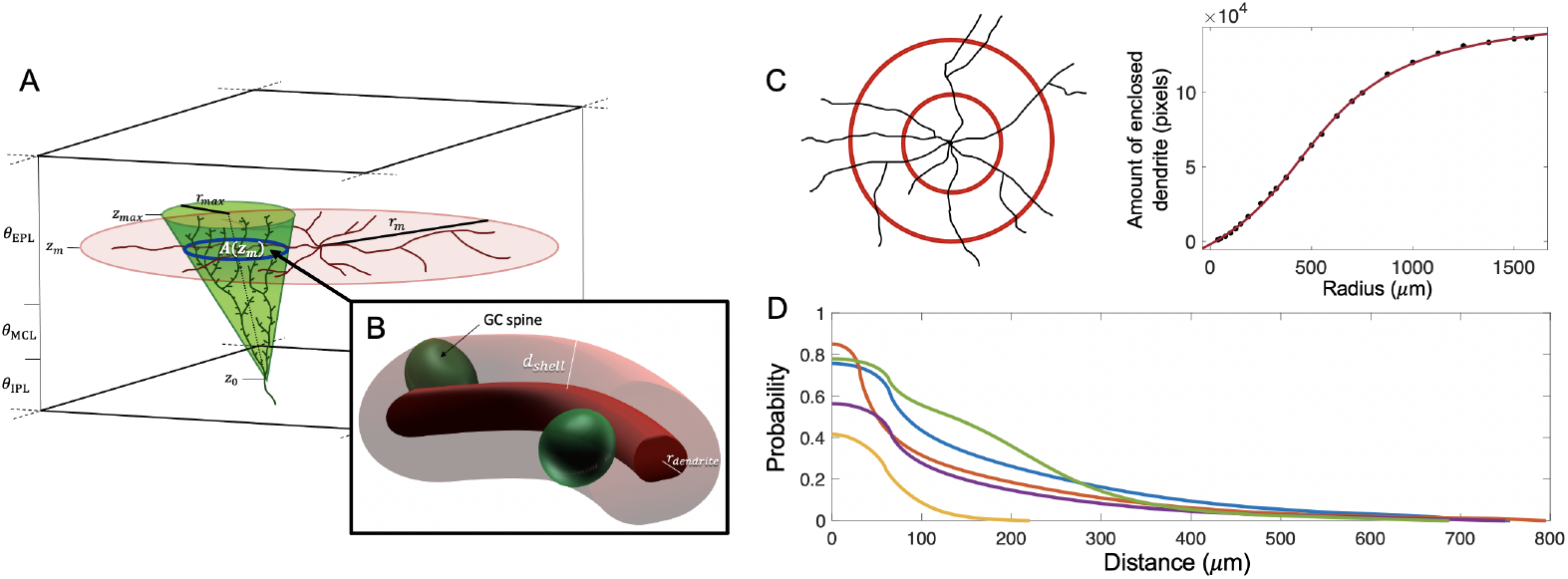
Schematic of the model. (**A**) Our model olfactory bulb (OB) has three layers: the external plexiform layer (EPL), mitral cell layer (MCL), and internal plexiform layer (IPL), each with a specified thickness. We modeled mitral cells (MCs) as flat disks with radius *r*_*m*_ (indicated in red), placed at at a height *z*_*m*_ in the EPL. We modeled granule cells (GCs) as inverted oblique circular cones (indicated in green), with bottom vertex at *z*_0_ in the MCL or IPL, and top face at *z*_max_ in the top half of the EPL; the radius of the top face was *r*_max_. By integrating the MC lateral dendrite density *ρ*_*m*_(*r*) over the area of intersection *A*(*z*_*m*_) between the MC disk and GC cone, we calculated the length of MC lateral dendrite contained in the overlap. (**B**) By treating the lateral dendrites as cylinders, we calculated the potential volume of interaction between MC lateral dendrite and GC spines around the length of MC lateral dendrite in the overlap. Presuming that GC spines are roughly evenly distributed at any given height along the cone, such that the spine density *ρ*_*g*_ depends only on *z*, and that this density is roughly constant, the expected number of spines in the interaction volume (which we treat as equivalent to the expected number of synapses) was *ρ*_*g*_(*z*) multiplied by the volume. (**C**) We utilized *camera lucida* images of MCs to calculate the total amount of dendrite within circles of increasing radii and found that this quantity was well fit by an equation *α* tan^*−*1^(*kr* + *β*) + *C* (mock MC shown). The equation for the density of lateral dendrites *ρ*_*m*_(*r*) was derived from this. For example, a cell image from [41] was well fit by the function with *α* = 66, 270, *k* = 0.002609, − *β* = 1.159, and *C* = 55, 010 (*r*^2^ value = 0.9998). (**D**) Sample probability curves for different MC/GC pairs, using the expected number of synapses as the mean of a Poisson distribution.

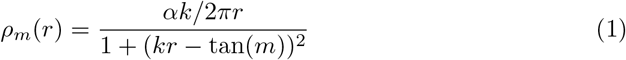

where *α, k*, and *m* are constants, and *ρ*_*m*_ lies between 0 and some maximum radius *r*_max_ (derivation in Methods Sec. 0.1).

GC dendrites pass roughly orthogonally to MC lateral dendrites, with the effective radius of the dendritic tree increasing along the height of the tree [8, 9, 40, 41]. We therefore defined the GC dendritic tree over an inverted oblique circular cone, with face parallel to the faces of the OB space. The volumetric spine density (number of spines per *µ*m^3^) at a given height *z* on the cone is calculated from dividing the vertical spine density *N*_*s*_ (defined as the number of spines per *µ*m height) by the cross-sectional area of the cone at *z* (Eq 33):

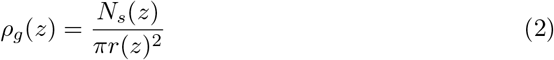

where *ρ*_*g*_ is defined over a range between some minimum and maximum heights *z*_0_ and *z*_max_ marking the bottom and top of the cone respectively. Since the vertical spine density of GCs tends to have an overall concave shape [8, 40], we modeled our vertical spine density as a simple parabola following [8, 40]. *ρ*_*g*_(*z*) can then be expressed as:

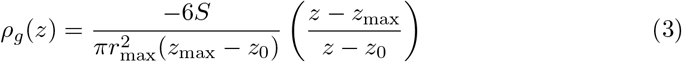

where *r*_max_ is the maximum cone radius and *S* is the total number of available spines on the GC (derivation in Methods Sec. 0.2).

We defined the average number of synapses between a MC-GC pair as the average number of GC spines within sufficient proximity to the lateral dendrites of an MC to establish synapses. Using the above distributions, we first calculated the length *L* of MC lateral dendrites present in the overlap between the MC and GC dendritic trees by integrating the MC dendritic density over the area of intersection between the two trees (Eq 39):

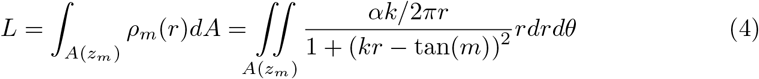

Here *A*(*z*_*m*_) is the area of intersection between the MC disk and the GC cone at the disk’s height *z*_*m*_. The integrals for the different cases, which depend on the relative position and sizes of the MC and GC, are shown in Methods, Section 0.3.

In order to account for the 3-dimensional nature of potential interactions between MC and GC, we converted the overlap dendritic length *L* into an equivalent volume by assuming the lateral dendrites to be cylinders of radius around 0.63 *µ*m [9]. We then defined the volume of interest to be the cylindrical sheath of thickness *d*_shell_ surrounding the lateral dendrite, with *d*_shell_ = 1.02 *µ*m, the effective diameter of a spine [59] (Fig 1B). The volume of this sheath was then:

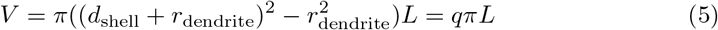

with *q* = 2.32 *µ*m^2^.

The density of GC spines in this volume was *ρ*_*g*_(*z*_*m*_), under the simplifying assumption that it was constant throughout the sheath volume. Thus, the expected number of spines in this volume, and by our definition synapses, was just:

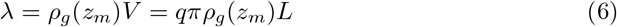

Because most MC-GC pairs make only one synapse [59], we used a Poisson distribution to calculate the probability of synapse from the expected number of synapses. If multiple synapses formed, they were treated as a single effective synapse. Thus, the probability of synapse was:

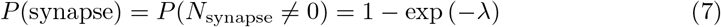

### Cell placement

To spatially distribute MCs and GCs, we modeled the OB as a thin circular cylinder with area *A* and thickness *θ* subdivided into parallel layers based on OB anatomy [60]. MC disks were distributed in the topmost layer of our model, the equivalent of the external plexiform layer (EPL), where interactions between GCs and MCs occur [2]. The number and location of these MCs was determined by their glomeruli, which are conglomerations of the apical dendrites of MCs. These glomeruli existed in the eponymous glomerular layer (not modeled) above the EPL, and we assumed that projections of these glomeruli onto the EPL were distributed randomly in the x-y plane, an easily relaxed assumption if future connectomic studies provide finer information. We calculated from data the overall area density of glomeruli in the OB (Methods, Sec. 0.3), so the number of glomeruli for a given OB space was just this density multiplied by the area *A*. Each glomerulus was randomly assigned between 15 to 25 MCs, whose disk centers were randomly scattered in the x-y plane around that glomerulus’s projection, with distance from the glomerular projection drawn from a truncated logistic distribution fit to data from [4] (Fig. S1). MCs were also divided into type I and type II subgroups (2:1 ratio), which differ in where in the EPL their lateral dendrites ramify [2, 40, 41]; thus the z-position of each MC in our model depended on its type assignation. The radius of an MC disk was drawn from a uniform distribution between 75 and 800 *µ*m, based on measurements of MC images from [9].

Once we placed the MCs, we added GCs. GCs can be divided into two major types based on where in the EPL their trees principally spread: the aptly named superficial and deep GCs, which are thought to interact primarily with tufted cels (TCs) and MCs, respectively [8, 61]. Since our excitatory population consisted of MCs, we only included deep GCs, although superficial GCs and TCs could easily be modeled too (see Discussion).

The vertices of the previously described GC cones were distributed uniformly randomly in the x-y plane of the OB space. We used images of GCs from [8] to roughly determine the bounds of GCs in the z-direction. The bottom vertex of the cone either inhabited the mitral cell layer (MCL), which lies directly below the EPL, or the internal plexiform layer (IPL), which is below the MCL and constituted the bottom-most layer of our model. Meanwhile, the top face of the cone was confined to the top half of the EPL, and the xy-position of its center was located a random distance away from the xy-position of the bottom vertex (*i*.*e*. potentially making the cone oblique).

For simplicity, we drew the total number of spines *S* for a particular GC from a uniform distribution determined by the cone’s volume. However, because only spines in the EPL are relevant for forming synapses with MCs, we limited the number of possible synapses each GC could make (*S*_available_) to the total number of spines present in the EPL, which could be found by integrating *N*_*s*_(*z*) from the bottom of the EPL to the maximum height of the cone:

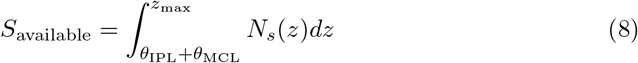

### Network generation

To generate the network, we added GCs individually and compared with each MC to determine, via the equations above, whether a synapse would exist between that pair. To account for preexisting synapses that an MC might already have (since spines corresponding to those synapses occupy space in the interaction volume surrounding the MC lateral dendrites), we weighted the calculated volume *V* = *qπL* for that MC-GC pair by the ratio of the volume of unoccupied interaction space to the volume of total interaction space on that MC, assuming for simplicity that the preexisting synapses are distributed evenly along the length of the dendrites:

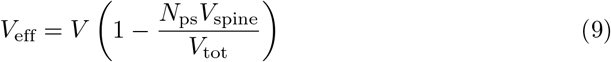

Here *N*_ps_ is the number of preexisting synapses, *V*_tot_ is the total volume of interaction space on the MC, and *V*_spine_ = 0.58 *µ*m^3^ is the average volume of a spine [59]. We used *V*_eff_ in Eq. 6 to determine the average number of spines and in turn the probability of synapse. We repeated this for each MC (whose order is shuffled for each new GC to avoid bias) until every MC in range has been tested. If the number of connections exceeded the maximum number of synapses allowed for that GC, we retained a random subset of those connections with size equal to the number of available synapses, and removed the remnant. Since the network generation was probabilistic, it was unlikely but not impossible that a GC would be disconnected from all mitral cells and thereby not contribute to the network. Thus, we generated GCs individually, discarding those that were disconnected from all mitral cells, until we reached a target number of GCs, which we calculate to be 15 deep GCs per MC based on current estimates of cell numbers in the OB [1]. Thus, we generated a model OB of radius 600 *µ*m (area 1.13 mm^2^) containing a network of 3,550 MCs and 53,250 GCs.

### Single cell dynamics

To explore network function we modeled individual MCs and GCs as dynamical systems described by the Izhikevich equations [42, 43]:

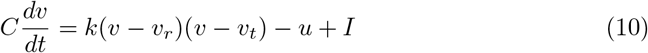

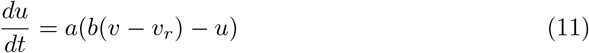

with spike reset:

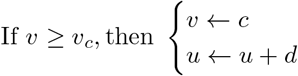

where *v* is the membrane potential; *u* is a recovery current; *v*_*r*_ is the resting potential; *v*_*t*_ is a threshold; *v*_*c*_ is a cutoff; *I* is an external current; and *a, b, c, d*, and *k* are free parameters.

We selected parameters to model class II behavior of MCs [62] and to establish realistic *f* − *I* curves [9] (Fig. 2C). Following conductance based models [5, 7], we took GCs to be integrators [42, 43]. Other work suggests that some GCs may display resonator properties, including subthreshold membrane potential oscillations [63, 64], but the very low oscillation frequency may make them irrelevant to excitability classification [43], especially since some GCs appear to be entirely non-resonant [63]. So, *b < a* in the Izhikevich model; additionally, by assuming *b* to be negative, we could take advantage of the following equations to calculate *b* and *k* [43]:

**Fig 2.**
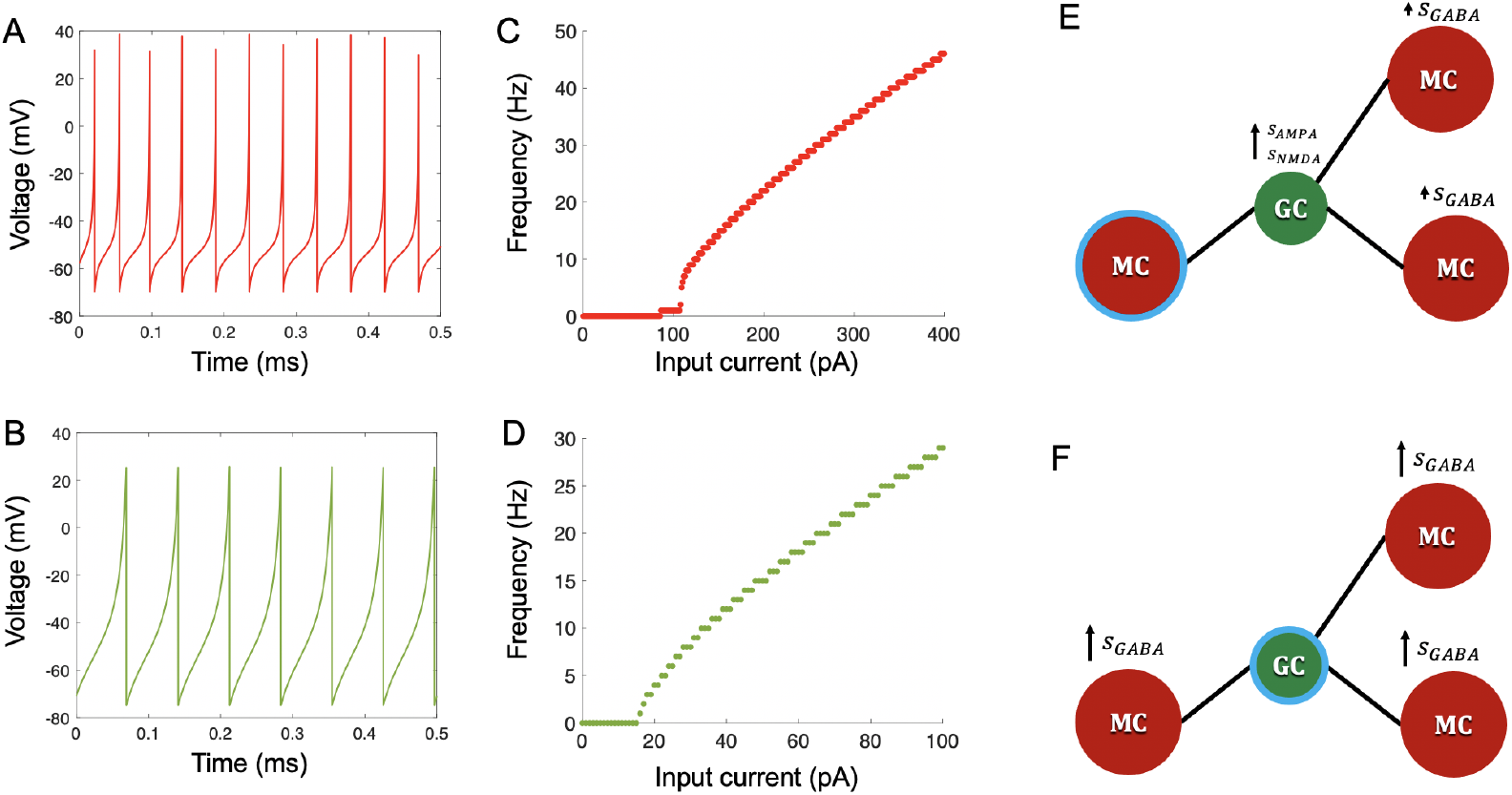
Cell dynamics modeling. (**A**) Sample voltage trace of an MC. Input was 200 pA direct current. (**B**) Sample voltage trace of a GC. Input was 45 pA direct current (**C**) Frequency-current (f-I) curve for an MC. (**D**) f-I curve for a GC. Each cell in (**C**) and (**D**) was simulated with the given parameters in Table 1, receiving direct current for 1 second of in-simulation time per current intensity. (**E**) When an MC spikes (MC with blue ring), gating variables of GC AMPA and NMDA receptors increase at synapses between that MC and all connected GCs. In turn, gating variables of MC GABA receptors increase by a smaller amount at synapses between connected GCs and their subsequent connected MCs (**F**) When a GC spikes (GC with blue ring), gating variables of MC GABA receptors increase at synapses between that GC and all connected MCs.

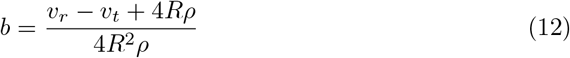

**Table 1.** Parameters for single cell internal dynamics

| Variable | Mitral cell | Granule cell |
| --- | --- | --- |
| $k$ (nS(mV)<sup>-1</sup>) | 2.5 | 0.067 |
| $a$ (ms<sup>-1</sup>) | 0.02 | 0.01 |
| $b$ (nS) | 12 | -0.133 |
| $c$ (mV) | -70 | -75 |
| $d$ (pA) | 13 | 2 |
| $v_r$ (mV) | -58 | -71 |
| $v_t$ (mV) | -49 | -39 |
| $v_c$ (mV) | 30 | 25 |
| $C$ (pF) | 191 | 48 |

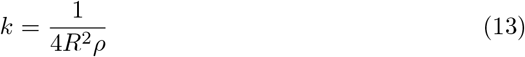

where *R* is input resistance and *ρ* is the rheobase (minimum DC current to produce spikes). We chose the remaining parameters to match a realistic *f* − *I* curve from [8] (Fig. 2D). The parameter values are given in Table 1. We drew parameters from a normal distribution with standard deviations equal to 1/10 of each mean, except for *b* and *k* for GCs, which we drew from normal distributions with standard deviations equal to 2/3 of the mean to satisfy the constraint that *b <* 0 while achieving a range of rheobases between 10 and 70 pA, and input resistances between 0.25 and 1.5 GΩ.

### Synaptic dynamics

We modeled dendrodendritic synapses for MC-GC pairs as NMDA and AMPA receptors on GCs, and GABA receptors on MCs. The synaptic AMPA current was

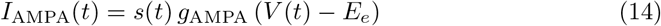

where *g*_AMPA_ is the conductance, *s*(*t*) is a gating variable representing the fraction of open channels, *V* (*t*) is the voltage of the recipient cell, and *E*_*e*_ = 0 mV is the excitatory reversal potential. For GABA receptors, we also noted that inhibitory signals from the cell periphery degrade as they propagate to the soma [65], which we described as an exponential decay. Including this decay, we modeled the GABA current as

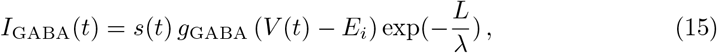

where *E*_*i*_ = − 70 mV is the inhibitory reversal potential, *L* is the distance between the MC center and the synapse, *λ* is a length constant, and other parameters were as for AMPA. The network generation did not identify MC-GC synapsse locations, so we chose points sampled randomly from the overlap between each MC and GC.

For NMDA receptors, we used [66]:

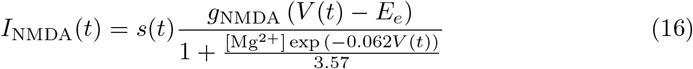

where the additional term in the denominator describes the magnesium block, with [Mg^2+^] assumed to be 1 mM [67].

The AMPA and GABA gating variables evolved as [68]:

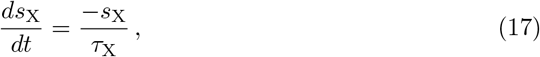

where *X* = GABA or AMPA. The NMDA dynamics followed

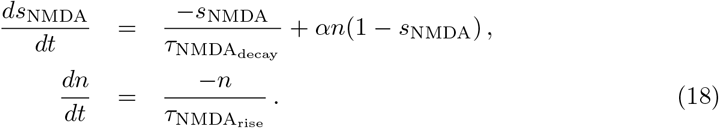

MC activation at reciprocal MC-GC synapses causes excitatory glutamate release onto NMDA and AMPA receptors on GCs, while GC activation causes inhibitory GABA release onto GABA receptors on MCs [69, 70]. Thus, when an MC spiked, the gating variables of GC NMDA and AMPA receptors at its synapses were updated as (Fig. 2E, center GC):

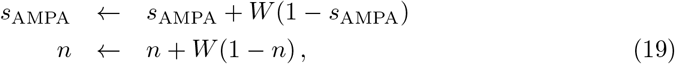

where *W* = 0.5. To account for network-driven GC activity [6], GABA gating variables of MCs indirectly connected to a spiking MC via shared GCs were also updated:

(Fig. 2E, right):

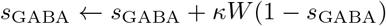

where 0 *< κ <* 1. When a GC spiked, the gating variables of MC GABA receptors at its synapses were updated as (Fig. 2F):

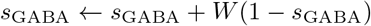

For each cell, the synaptic input at any time was the sum of the currents for its receptors at all synapses (NMDA and AMPA for GCs, GABA for MCs). We derived time constants *τ* and *α* from data [69], and tuned conductances and *κ* to reproduce lateral inhibition results from [71] as faithfully as possible. We calculated the length constant *λ* from the formula in [65] for the diameter of dendrite used in the connectivity algorithm (*d* = 1.26 *µ*m). Parameters are in Table 2.

**Table 2.** Synaptic parameters

| Parameter | Value |
| --- | --- |
| $g_{\text{AMPA}}$ (nS) | 0.73 |
| $g_{\text{NMDA}}$ (nS) | 0.84 |
| $g_{\text{GABA}}$ (nS) | 0.13 |
| $\kappa$ | 0.006 |
| $\tau_{\text{AMPA}}$ (ms) | 5.5 |
| $\tau_{\text{NMDA}_{\text{rise}}}$ (ms) | 10 |
| $\tau_{\text{NMDA}_{\text{decay}}}$ (ms) | 80 |
| $\tau_{\text{GABA}}$ (ms) | 18 |
| $\lambda$ ($\mu\text{m}$) | 675 |
| $\alpha$ ($\text{ms}^{-1}$) | 0.1 |

### Sister mitral cells are weakly correlated in the network

We asked what our local connection rules predicted for global network features such as MC to GC connectivity and vice versa. The distribution of MC connectivity to GCs was well fit by an exponential (Fig. 3A, top), as were individual type I and type II distributions (Fig. 3A, bottom). However, type I MCs connected to more GCs than their type II counterparts, likely because type II MC dendrites ramify higher in the EPL and thus overlap less with deep GC dendritic baskets. The distribution of GC connectivity to MCs was well fit by a skewed normal distribution (Fig. 3B; see Methods). These structural predictions can be tested as bulb connectomes become available. We also found that sister MCs, i.e., MCs connected to the same glomerulus, connected to more of the same GCs than non-sister MCs, but, surprisingly, this overlap was low (mean = 0.13) (Fig. 3C). This predicts that sister MCs, despite originating in the same glomerulus, will have distinct lateral dendrite synaptic patterns consistent with [4], and will encode odors non-redundantly, perhaps explaining findings in [3, 72].

**Fig 3.**
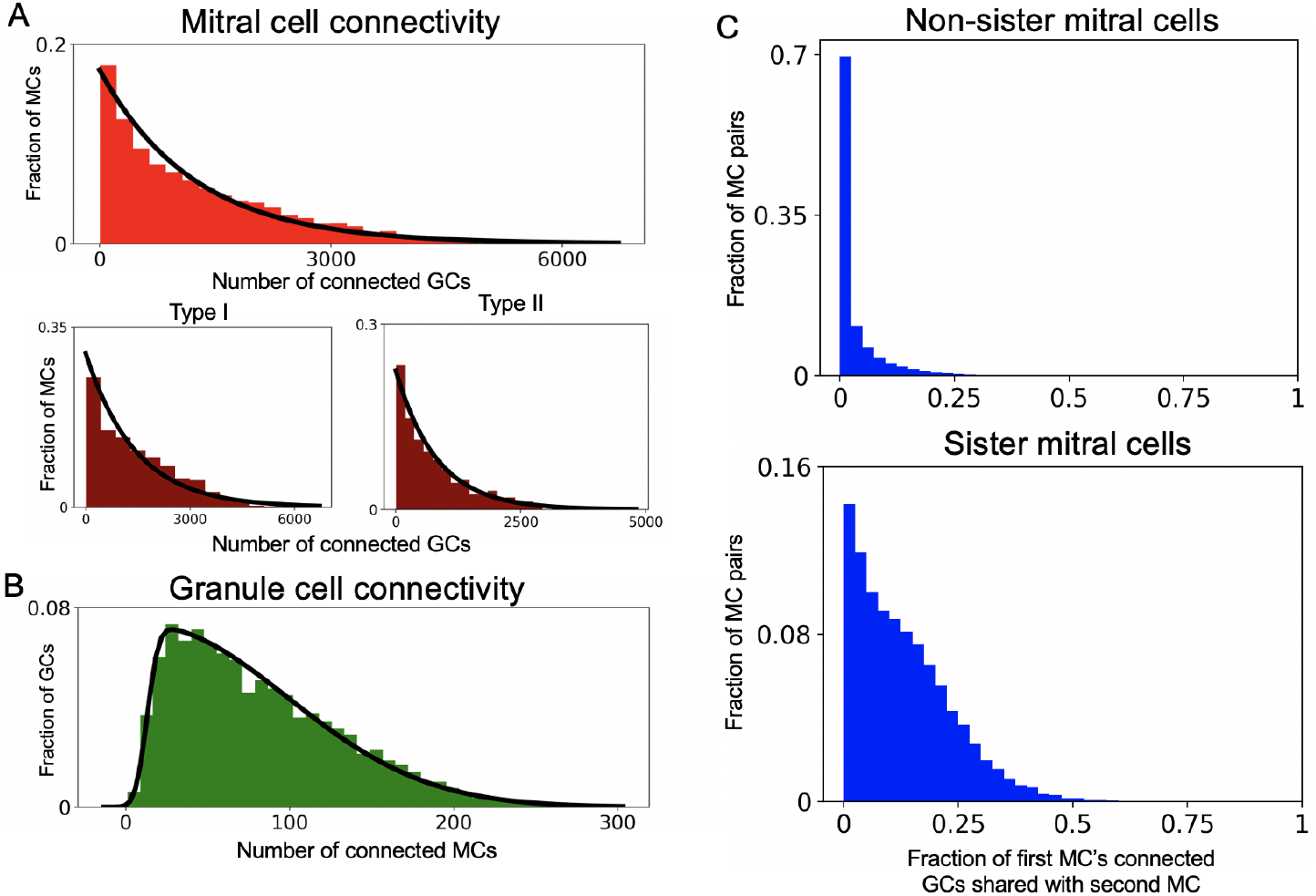
Network connectivity. (A) Top: Distribution of MCs by number of connected GCs, black line = exponential fit (mean 1225.8; *χ*^2^ = 124.52, *p <* 10^*−*14^). Bottom: Distribution of type I (left) and II (right) MCs by number of connected GCS. Black line for type I MCs = exponential fit (mean 1426.0; *χ*^2^ = 139.1, *p <* 10^*−*22^). Black line for type II MCs = exponential fit (mean 818.7; *χ*^2^ = 39.9, *p* = 0.022). (B) Distribution of GCs by number of connected MCs. Black line = skewed normal fit (parameters *α* = 15.2, *ξ* = 13.0, *ω* = 85.5; *χ*^2^ = 501.1, *p <* 10^*−*83^; see Methods for definition of skewed normal) (C) Distribution of fraction of connected GCs shared with another MC for non-sister (top) and sister (bottom) MCs.

### Network oscillations and granule cell inhibition

There are prominent local field potential (LFP) oscillations in the OB, with different frequencies associated to specific aspects of olfaction, e.g., fine odor discrimination and associative learning [55]. Thus, we tested whether respiratory baseline and odor input produced oscillations and in what range. MCs received Poisson inputs with time-varying rates [73] from olfactory sensory neurons (OSNs) activating 100 synapses per MC, each with an NMDA and AMPA receptor modeled as Eqs. (17-18), albeit with conductances and time constants based on [74] (Table 3). For each synapse, a spike input caused the NMDA and AMPA gating variables to increase as in (19).

**Table 3.** External input parameters

| Parameter | Value |
| --- | --- |
| $g_{\text{AMPA}}$ (nS) | 6.7 |
| $g_{\text{NMDA}}$ (nS) | 12 |
| $\tau_{\text{AMPA}}$ (ms) | 14.3 |
| $\tau_{\text{NMDA}_{\text{rise}}}$ (ms) | 13 |
| $\tau_{\text{NMDA}_{\text{decay}}}$ (ms) | 70 |
| $\alpha$ (ms<sup>-1</sup>) | 0.03 |

To simulate respiratory baseline, we modeled the Poisson rate *r*(*t*) for the external inputs as

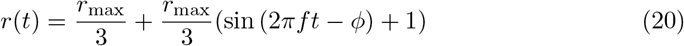

with *f* = 2 Hz, representing respiratory rate. To determine *r*_max_ we drew *x*_*g*_ uniformly from 0 to 0.25 Hz for each glomerulus; then for each sister MC of this glomerulus, we drew *r*_max_ from a Gaussian (mean *x*_*g*_, standard deviation *x*_*g*_*/*10). Similarly, to determine the phase *ϕ*, we first drew *p*_*g*_ uniformly from 0 to 2*π* for each glomerulus since phases of non-sister MCs are highly uncorrelated [72]. Then for each MC in a glomerulus we drew *ϕ* from a Gaussian (mean *p*_*g*_, standard deviation *π/*4) reflecting variability among sister MCs [72].

To simulate odor input, we increased respiratory rate to 6 Hz to represent sniffing and normalized *r*(*t*) to be stronger:

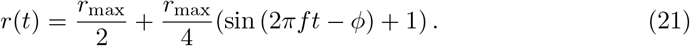

We selected 0.2 of the glomeruli to receive odor input and resampled *r*_max_ for all MCs, except that for odor-receiving glomeruli, *x*_*g*_ was drawn uniformly between 2 and 3 Hz, while for non-odor-receiving glomeruli, *x*_*g*_ was drawn uniformly between 0 and 0.25 Hz; *ϕ* was resampled as before.

We used [75, 76] to calculate LFPs:

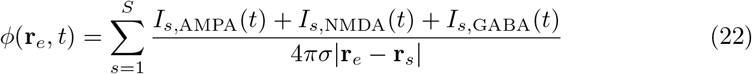

where for each synapse *s*, **r**_*s*_ is the location, *I*_*s,X*_ (*t*) is the current through a receptor type, and *σ* is extracellular conductivity (1*/*300 Ω^*−*1^ cm^*−*1^). Here, **r**_*e*_ is the “electrode” location at the xy-center and halfway up the EPL. After filtering and selecting the region of interest, we acquired the LFP power spectrum averaged across 10 trials (Methods; Fig. 4).

**Fig 4.**
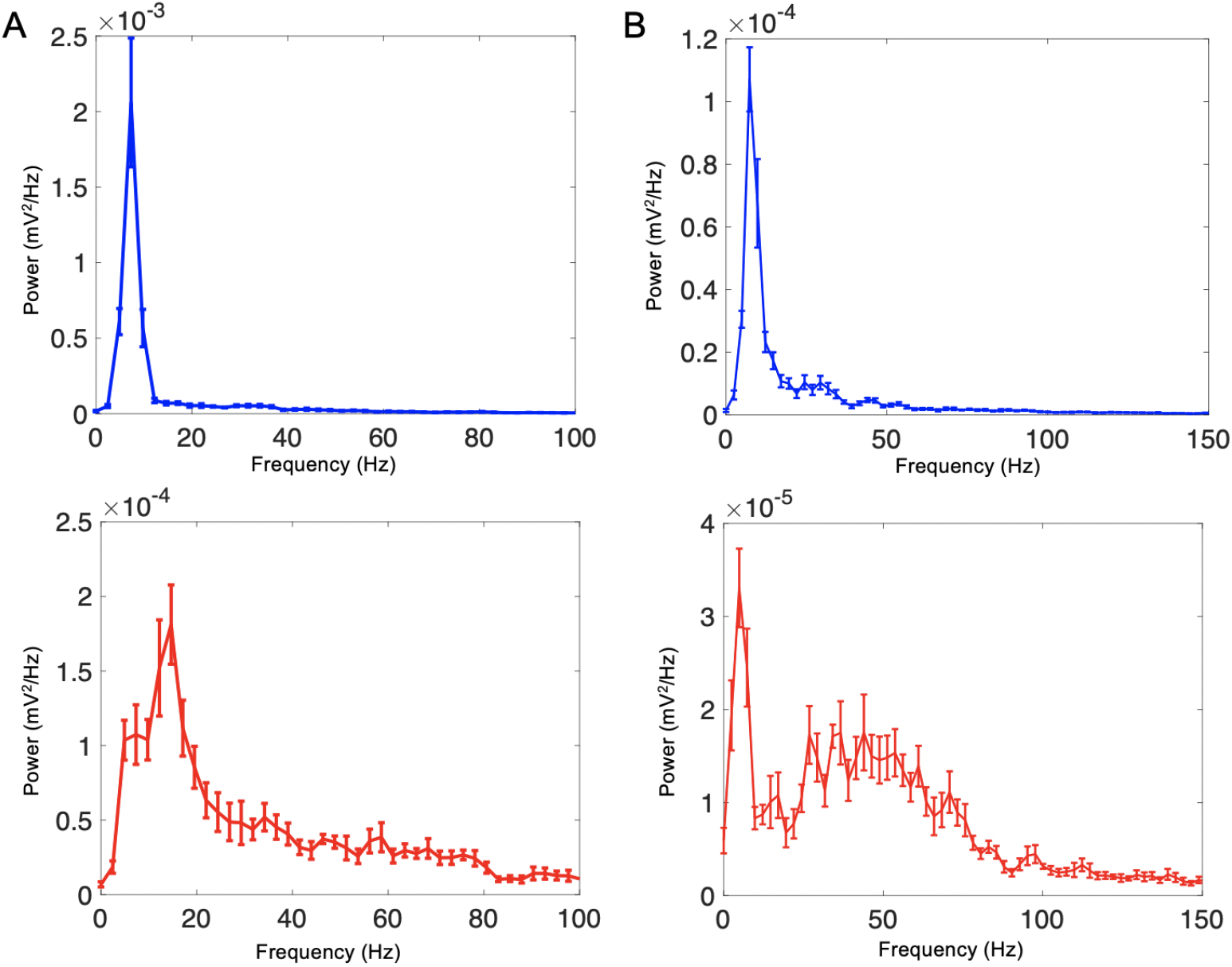
Oscillatory dynamics. (A) A network with a 15:1 GC:MC ratio and respiratory input exhibited LFP oscillations at 7 Hz in the theta range (2-12 Hz) (Top). Odor input to a subset of glomeruli, caused oscillations at ∼ 15 Hz, in the lower beta range (15-40 Hz), in addition to activity at 6 Hz, corresponding to respiration (Bottom). (B) A network simulating tonically inhibited GCs, and hence a lower 5:1 active GC:MC ratio. Respiratory input exhibited LFP oscillations at 7 Hz in the theta range (Top). Odor input additionally caused oscillations with a broad frequency peak around 40-55 Hz, in the gamma range (35-100 Hz) (Bottom). Curves averaged from 10 trials; bars indicate standard error.

During baseline respiration, the LFP oscillated at 7 Hz, in the theta range, consistent with data showing theta oscillations coupled to resting respiration [55] (Fig. 4A, top). During odor input, the LFP oscillated at 6 Hz, corresponding to sniff rate, and at 15 Hz, in the beta range (Fig. 4A, bottom). This was surprising, since odor presentation is thought to produce gamma oscillations (35-100 Hz) due to activity at the MC-GC synapse, while beta oscillations are more commonly associated with cortical feedback to the OB [14, 55]. However, given that a major role of cortical feedback is to activate GCs [77], and that without such feedback, large segments of GCs are tonically inactivated [1, 38, 78], we hypothesized that reducing the number of active GCs to approximate tonic inhibition might produce gamma oscillations, especially since other computational studies demonstrating gamma have utilized lower GC:MC ratios [11, 14]. Therefore we repeated the experiment with a GC:MC ratio set to 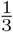 that of the full network. Indeed, odor presentation led to LFP oscillations peaking around 40–55 Hz, in the gamma range (Fig. 4B). This suggests that overall activity of the GC network, determined by a balance between tonic inhibition and excitatory feedback, is a major determinant of whether the OB oscillates in the beta or gamma ranges during odor presentation [79, 80]. This is in line with studies demonstrating the importance of GC excitability to LFP oscillation frequency [14, 23, 39].

### Lateral inhibition follows network architecture

It is believed that lateral inhibition by granule cells may be involved in gain control, synchronization of MC output, and decorrelation of odor representations [34, 71, 81–86]. To ask how recurrent interaction between MCs and GCs varied with distance, we first measured the number of GCs shared between pairs of MCs at around the same height in the EPL, and connected individually to similar number of GCs. This shared number decreased with distance following a relation of the form *a* exp(− *bx*^*n*^) (Fig. 5A), with *n* ∼ 1.5 intermediate between an exponential and a Gaussian, leading us to anticipate that lateral inhibition between pairs should be of similarly short range. To test, we selected MC pairs as above, and mimicked the experimental methodology of [71], where we measured the firing rate of one MC when (a) it alone was excited with direct current and (b) when the other MC of the pair was also excited. We found that the decrease in firing rate of the first cell in conditions (a) vs. (b) declined with separation between the pair and had a similar short-range form to the relationship between MC separation and number of shared GCs (Fig. 5B). For comparison, we built a second network with the same number of cells and average MC-GC connectivity but with constant, distance-independent probability of connection for each MC-GC pair. In this second network, both the number of shared GCs and magnitude of lateral inhibition were constant and independent of distance (Fig. S2). This suggests that the assumption of random connectivity used for simplicity in many studies [10, 20, 21, 23] leads to fundamentally different inhibitory effects in the feedforward olfactory pathway.

**Fig 5.**
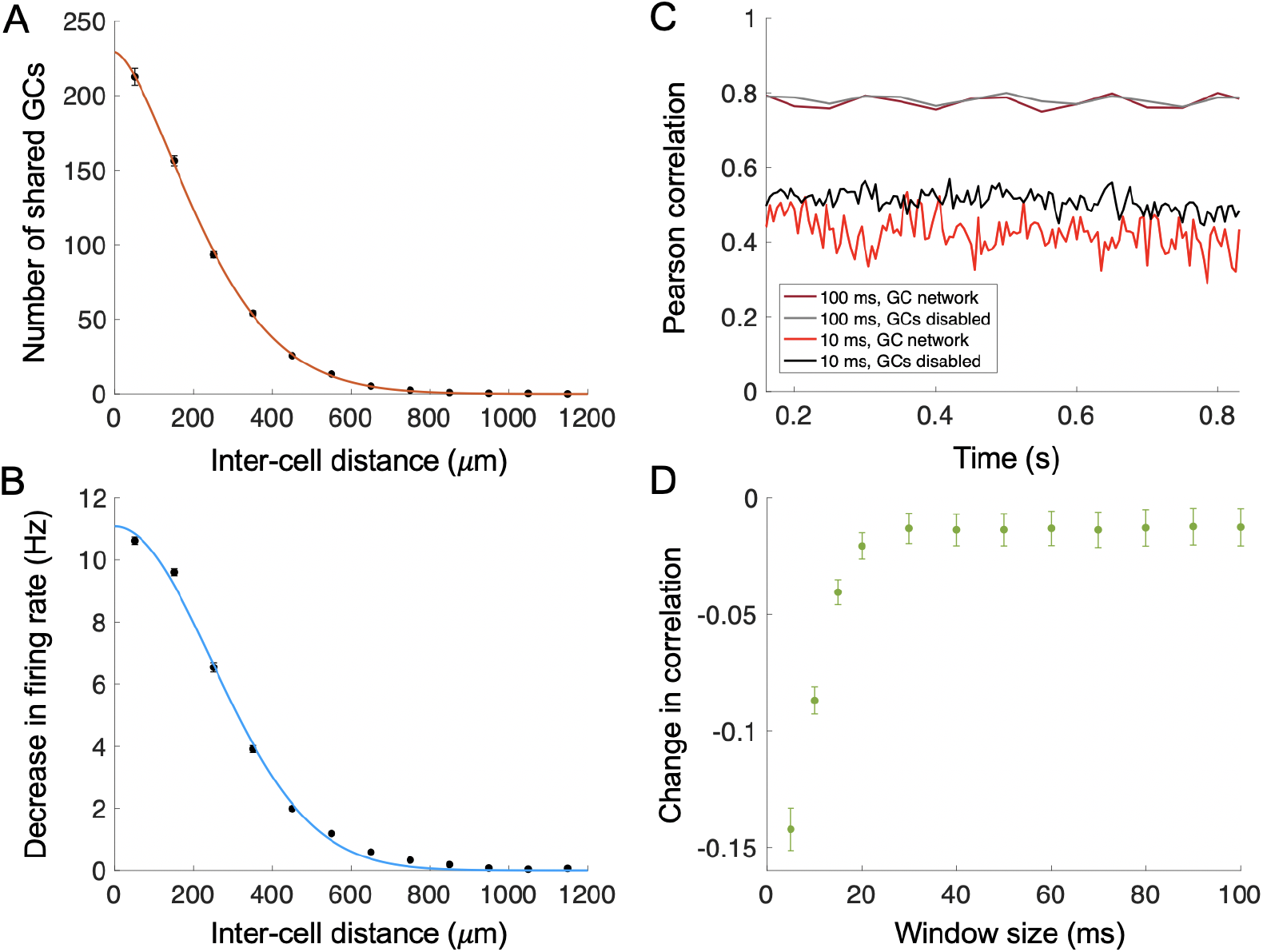
Network connectivity, lateral inhibition, and decorrelation. (A) The number of shared GCs between pairs of MCs (*n* = 1436). MC pairs were partitioned based on inter-cell distance into 12 bins each representing a 100 *µ*m span. Each point represents the average number of shared GCs of all MC pairs in a given bin and was placed halfway along the bin width, with the bar representing standard error. The red curve is a fit of the form *a* exp(−*bx*^*n*^), with *x* in microns, *a* = 229.2, *b* = 1.721e-04 and *n* = 1.545. (B) Lateral inhibition strength decreases with distance between MCs. MC pairs (*n* = 1436) were simulated with one and then both cells receiving current input, and the resulting decrease in firing rate for the first cell was measured. MC pairs were again partitioned based on inter-cell distance into 12 bins each representing a 100 *µ*m span. Each point represents the average decrease in firing rate of all MC pairs in a given bin and was placed halfway along the bin width, with the bar representing standard error. The blue curve represents a fit of the form *a* exp(−*bx*^*n*^), with *a* = 11.08, *b* = 9.375e-06 and *n* = 1.976. (C) Example of evolution of Pearson correlation between model OB responses to two odors, each targeting 30 of 178 glomeruli (25 overlapping) over time, in the presence and absence of GC activity. Dark red/grey: GCs enabled/disabled for 100 ms time window. Bright red/black: GCs enabled/disabled for 10 ms time window. (D) The decrease in Pearson correlation was maximal for shorter timescales. We measured mean correlation over the period after the first sniff (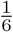 of a second) for networks with and without GCs, and then calculated the difference of the two. The magnitude of decorrelation increased as the window size decreased below 20 ms. Data averaged over *n* = 15 odor pairs, each odor activating 30 glomeruli, and with varying overlap ranging from 5 to 25 glomeruli (average overlap = 18). Error bars = standard error of the mean.

Given that the magnitude of the decrease in firing rate for an MC produced through lateral inhibition was relatively low (Fig. 5B) compared to the firing rate produced through direct current stimulation of the MC alone (∼70 Hz), we were curious whether the GC network in our model could effectively decorrelate odor patterns. To answer this question, we generated a set of 6 odors, with odor 1 targeting glomeruli 1 through 30, odor 2 targeting 6 through 35, and so on, leading to an average overlap between odors of ∼18 glomeruli; we then simulated our system receiving each of these odors. We ensured that odor inputs for the overlapping glomeruli were delivered at the same phase and strength, because input phase differences can already decorrelate responses, and we were primarily interested in the specific role played by the GC network. We measured the Pearson correlation of the MC firing rates induced by each odor within sliding time windows of fixed duration. Afterward, we repeated the same experiment, except with the GC network disabled by setting the GABA conductances on MCs to zero (Fig. 5C). Then, for each condition we time-averaged the correlation after the first sniff (by which point the system had equilibriated). Finally, we calculated the difference between the odor-response correlations measured with and without the granule cell network, and treated this as the amount of decorrelation induced by the GC network. Without granule cells (black and grey lines in Fig. 5C), we found that the MC output correlation reflected the number of overlapping glomeruli when measured over long time windows, but was less correlated over short windows because of variations in spike timing for individual MCs determined by the dynamics. At short timescales, we observed that the GC network had a small but significant decorrelating effect, especially when the response correlation was measured in windows of *<*20 ms (Fig. 5C,D). This is consistent with experiments showing that GCs primarily operate along such timescales and produce decorrelation through alteration of spike timing rather than gain control [87].

### Network architecture shapes cortical feedback

GCs and MCs receive extensive cortical feedback, which plays a major role in shaping odor representation as information is conveyed through the OB to cortex [28, 77, 88–92]. We thus asked how the arrangement of GCs in the network determined the expression of this feedback in the OB. We first examined the effect of external activation of the GC network on odor-receiving MC output, and how this effect varied with the spatial pattern of GC activation. Thus, we targeted excitatory feedback randomly to between 0.1%-20% of all GCs during presentation of an odor, and then compared to simulations where no feedback was present. The first two trials used the same network but targeted non-overlapping sets of GCs, in order to ascertain whether the effect of feedback could be attributed to which set of GCs was targeted. In the third trial, we performed the same experiment but in a different network which had the same arrangement of MCs but a different configuration of GCs. We calculated the change in odor-receiving MC firing rates with feedback for each trial and then computed the correlation of these changes over time between the first and second trials (same GC network) and between the first and third trials (different GC networks) using a sliding window of length 10 ms and 50% overlap between windows.

To our surprise, the particular arrangement of the feedback in terms of which GCs were selected had little effect on which MCs were ultimately affected, since the average correlation between trials in the same GC network (Fig. 6A, purple) was relatively high even for small amounts of targeted GCs. Moreover, the average correlation values between trials in different GC networks was significantly lower in comparison (Fig. 6A, blue). Thus, our results suggest that a primary determinant of the effect of cortical feedback on the bulb output is the network architecture of GCs, as opposed to which of these cells are specifically targeted. This implies that theories of feedback and neurogenesis in the bulb that rely on specific targeting of GCs and MCs [12, 28, 93] may require additional components beyond the basic MC-GC network to be feasible.

**Fig 6.**
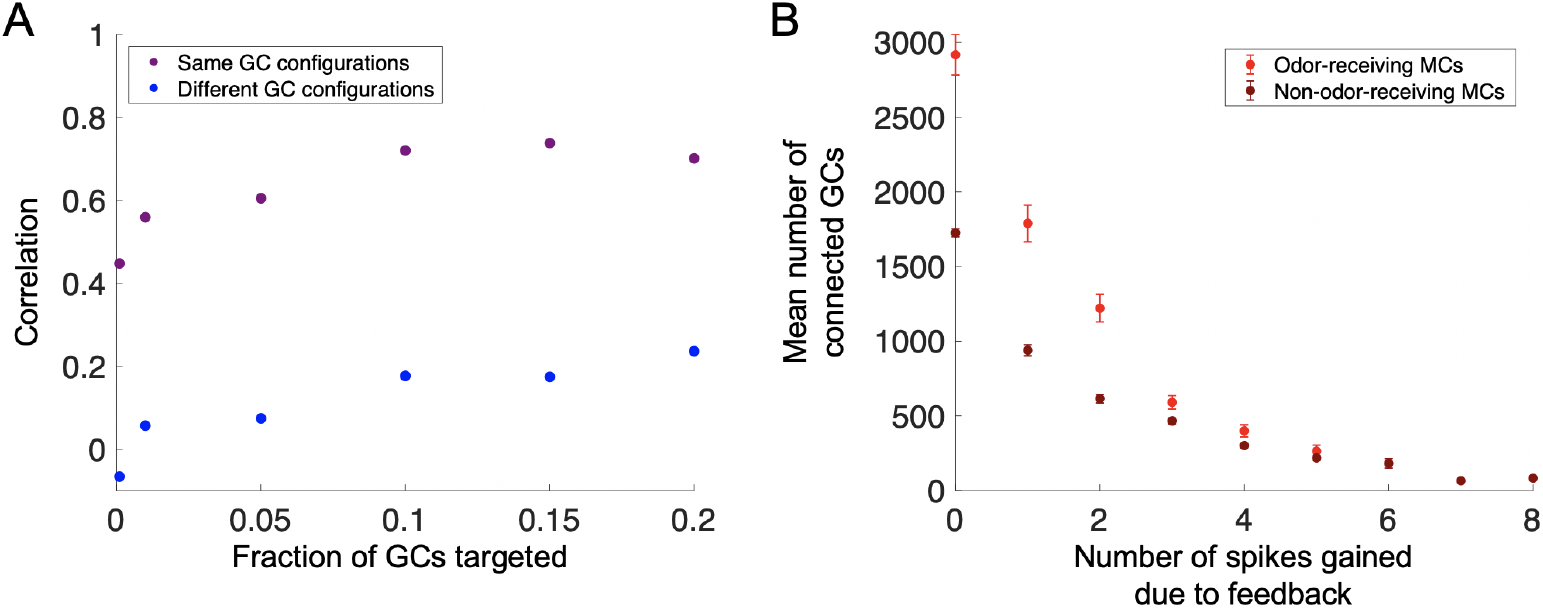
Network architecture and effects of external feedback to the OB. (A) The effects of randomly distributed, excitatory feedback to GCs depend on the OB network’s inherent connectivity and are relatively invariant to the specific GCs targeted. We presented an odor to the network without and then with external feedback to GCs, and measured the change in MC firing rate. For each feedback level, we ran three trials; the first two took place in the same MC-GC network, but targeted non-overlapping sets of targeted GCs. The third trial took place in a second network with the same MCs but a different spatial configuration of GCs. We computed the Pearson correlation for successive time windows of length 10 ms between the vector of feedback-driven changes in odor-receiving MC firing rates for the first two trials, in the same network (purple), and the first and third trial, in different networks (blue). We found that even for small numbers of targeted GCs, the mean correlation over time was high for the same-network trials despite targeting different sets of GCs, but was lower for the different-network trials. (B) MCs which connect to the least GCs are most affected by direct excitatory feedback. We randomly distributed excitatory feedback to 20% of MCs during odor presentation and recorded the resulting change in firing rate. MCs with larger firing rate increases tended to connect to a smaller number of GCs for both odor-receiving (red) and non-odor-receiving (dark red) MCs. Vertical lines indicate standard error. Data compiled from *n* = 5 trials.

We also examined how changes in MC firing rate due to direct positive feedback depended on the arrangement of GCs in the OB [94, 95], by first presenting the OB with an odor, and then presenting the same odor with excicatory feedback current to a randomly selected set of MCs. Repeating this experiment for different odors, we found that cells whose firing rates increased the most with feedback were connected to the least number of GCs, both for odor-receiving (Fig. 6B, red) and non-odor-receiving MCs (Fig. 6B, dark red). Thus, just as with GC feedback, the effect of direct MC feedback also appears to depend on the particular architecture of the GC network around MCs.

This model also supports findings on feedback-driven odor discrimination and generalization from a statistical model presented in [53].

## Discussion

The functions of the olfactory bulb, which reshape odor representations before they are sent to cortex, emerge from the complex dynamics of a structured network of granule cells, mitral cells and other cell types. Computational models are necessary for determining these collective dynamics, but are challenging to manipulate because of the size and complexity of the network. Many models navigate this challenge by building networks with a reduced number of neurons or GC:MC ratio [10, 11, 13, 14, 19, 22, 23] and either random connectivity or simple distance-dependent functions to generate connectivity [10–22]. These works have replicated several experimentally observed phenomena, such as LFP oscillations [11, 13–15, 17, 20, 23], odor decorrelation [21, 26, 83] and generalization [93], and even odor discrimination in complicated environments [12]; moreover, they have offered potential mechanisms for the relationship between cortex and OB via processes such as feedback and neurogenesis [12, 25, 26, 28, 47, 93].

However, the functional consequences of the anatomical constraints imposed by the OB’s cellular morphology remain unclear. Thus, we developed a modeling approach that employed geometry and dynamical systems theory to integrate realistic details of single cell anatomy [2, 8, 9,40, 41,61] and physiology [8, 9, 62, 71], along with empirical information about synaptic architecture [59], into a tractable, yet realistic, computational model of the OB network. The model simulates the activity of tens of thousands of cells on a standard laptop, and can be parallelized for greater speed. A key advantage of the model is its extensibility: additional cells and their connections could be added as shown here for MCs and GCs. This ease of making alterations to the network may be of use, for example, in the study of GC neurogenesis, which is known to play a crucial role in olfactory learning [35,36, 96–99].

One study attending to anatomy in large-scale OB modeling is [100], whose authors simulated individual lateral dendrites for each MC along a curved model of the OB, a level of detail limiting the number of MCs to an order of magnitude fewer than in our model. Their connectivity distributions for MCs to GCs and vice versa were Gaussian, while ours are heavily skewed, with a low peak and a long tail (Fig. 3). We considered that this difference may arise because MCs residing near the edges of our flat OB have curtailed lateral dendritic fields and hence presumably form fewer connections with GCs, while the curved OB of [100] has a smaller edge for a given surface area, leading to less skew. However, we found that periodic boundary conditions, which lead to *no* curtailed dendritic fields, are still far from having Gaussian connectivity (Fig. S3). The key difference is likely that [100] assumed fixed MC-GC synapse density per length along the MC lateral dendrites. The normal distribution of the MC dendritic field in [100] then implies normally distributed connectivity. Our model includes an additional spatial constraint: MCs “compete” for GCs since GCs each have a limited number of spines. Thus MCs with the most favorable lateral dendritic distributions for a given GC configuration form the most synapses, leading to the exponential distribution for MC connectivity. Indeed, if we counter-factually assumed a fixed distance-independent probability of MC-GC synapses, our model also produced normal connectivity distributions (Fig. S2).

Functionally, these differences manifest in the effects of lateral inhibition. Our results show that, statistically, lateral inhibition is more likely to be stronger over shorter distances as a consequence of the shape of the network’s distance-dependent connectivity. In contrast, Migliore *et al* specifically demonstrate an example where MCs which are far apart have a significant influence on each other’s firing. The appearance of such long-range inhibition is unlikely in our anatomically grounded model which uses a lower GC:MC ratio of 15:1 (unlike ratios between 20:1 and 100:1 in [100]), and realistic statistics of dendritic connection (unlike fully connected dendrites in [100]). It is conceivable that the prior synaptic tuning performed in [100] to simulate odor learning may allow the sparser inhibitory connections in our model to have a stronger long-range effect. Since we perform no such tuning and presume all synaptic strengths to be of roughly equal magnitude, the network’s natural, distance-dependent architecture determines lateral inhibition. Thus, network remodeling, whether through synaptic plasticity [27] or neurogenesis [36] may be key to overriding the anatomical constraints which would otherwise disfavor long-range interactions. This explanation may also contribute to resolving conflicting reports of the extent of lateral inhibition in the mammalian OB, with some experimental studies reporting that interactions are at least partially short-range [101–104] consistent with our findings, and others reporting distance-independent inhibition [105–107].

Our model predicts that with a realistic OB network topology and single cell response physiology, gamma oscillations, classically associated with feedback-independent OB activity after odor input [55], only appear with a lower active GC:MC cell ratio than the anatomical proportion. At the full ratio, we see beta oscillations, more commonly associated with activation of cortical feedback to the bulb during odor input [55], an effect that we did not explicitly model. Our results are consistent with previous experiments and models [14, 23, 39] which suggest that gamma oscillations appear when GC activity is reduced due to a lower baseline excitability than suggested by single cell neurophysiology, perhaps due to the influence of centrifugal [108] or deep short-axon cell inhibition [109]. Alternatively, the absence of gamma at high active GC:MC ratio could reflect the possibility that GC activation generally produces oscillations predominantly in the beta range, with other types of EPL interneurons being responsible for gamma oscillations [1]. Additionally, GCs have local and spike-independent processes [6] which may play a role in gamma oscillations [14, 20]; our model utilizes point neurons in order to facilitate large-scale simulation and does not fully capture such processes. Future experimental and theoretical work can separate these possibilities

Our model also suggests that cortical feedback to the bulb will be heavily guided by the existing network structure. Specifically, we found that the MCs affected by feedback to GCs were largely determined by the network configuration, rather than by the GCs targeted. Likewise, the local connectivity to GCs determined which MCs responded most to direct cortical excitation. Similarly, we found that sister MCs originating in the same glomerulus exhibited highly non-overlapping connectivity patterns (Fig. 1C). This prediction is consistent with previous experimental studies [3, 4], but differs from many models, which, for simplicity, treat all MCs associated to a glomerulus as equivalent [15, 18, 21, 84, 110–113]. Thus, our results suggest the importance of accurately including network structure for determining the function of individual cells in the OB and their collective behavior, as well as the power of our model in providing a way to probe this relationship.

In the future, it will be interesting to extend our model by introducing other OB cell types, notably tufted cells (TCs) and their corresponding superficial granule cells. TCs form the second major population of excitatory neurons in the OB and have different anatomical and electrophysiological properties from MCs [2, 9, 40, 41]. Moreover, they may have significantly different functionality from MCs in odor processing [88, 114–116]. For their part, superficial GCs also have anatomical and electrophysiological properties which separate them from deep GCs [8, 40, 61]. Indeed, based upon their location in the EPL, they likely form connections with type II MCs in addition to TCs, and, in our model, their absence may potentially explain the relatively lower number of GCs connected to type II MCs as compared to type I MCs. Thus, the addition of such cell types to our model will facilitate exploration of the differences between MCs and TCs as well as between the two MC types. Perhaps such studies will shed light on a classic question: why do neural circuits like the OB need so many different cell types with different properties to carry out their functions, rather than simply having more complex circuitry connecting fewer functional types as in silicon hardware [117]?

## Methods

All values were derived, where possible, from measurements of the murine olfactory bulb [4,8, 9, 33, 118], the chief exception being the derivation of MC lateral dendritic density, which was based on images from study of rabbit [40] and rat [41].

### 0.1 Mitral cell lateral dendritic density

Analyzing *camera lucida* drawings of mitral cells from [40] and [41], we fit a function of the following form for the total dendritic length contained within a circle of radius *r*:

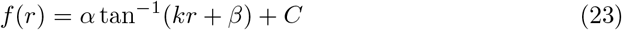

Then by default:

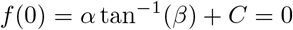

and:

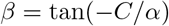

So Eq 23 becomes:

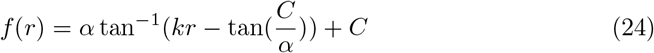

where *r* ∈ [0, *r*_max_] and 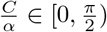 . Replacing for convenience *C* with *mα*, Eq 24 then becomes:

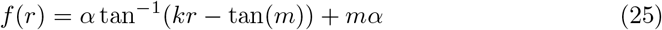

with 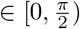

Now for an annulus of thickness *ϵ*, the ratio of the additional length of dendrite encapsulated in the annulus to the area of the whole annulus is:

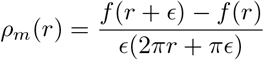

In the limit as *ϵ* goes to zero, we see that this equation simply becomes:

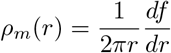

Plugging in *f* (*r*), Eq 25 for the dendritic density finally becomes Eq 1:

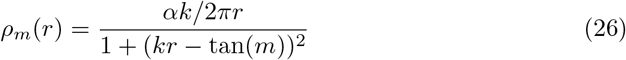

*α, k*, and *m* are defined in terms of the variables *γ, ξ*, and the maximum radius *r*_max_. *γ* ∽ Uniform(0.2, 0.3) describes the fraction of *r*_max_ where the maximum of 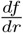 occurs. *ξ* ∽ Uniform(1*/*3, 4*/*5) represents the fraction of the maximum value of 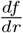 at *r* = 0. Additionally, we assume that the total length of dendrite *L* for a given MC is proportional to the area of that MC’s dendritic field, such that:

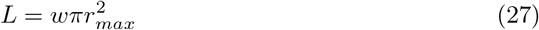

where we set *w* ∽ Uniform(0.00255, 0.00510) for each MC. Then, since the maximum value of 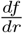 is *αk* and occurs at *r* = tan(*m*)*/k*, it follows that:

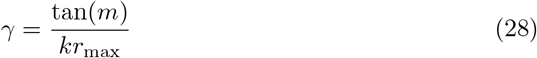

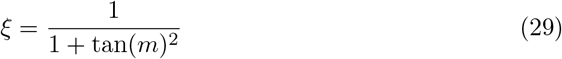

and the values of *m, k*, and *α* can be re-expressed as:

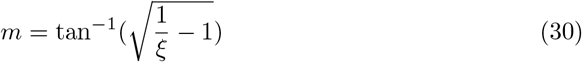

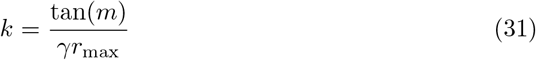

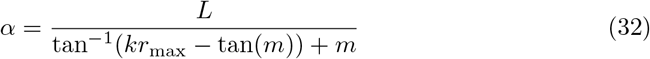

### 0.2 Granule cell spine density

The equation of spine density (in spines per unit volume) as defined in Eq 2 was:

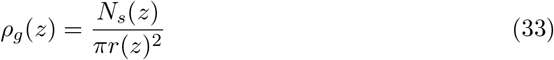

For *z* ∈ (*z*_0_, *z*_max_], where *z*_max_ and *z*_0_ are the maximum height and bottom of the dendritic tree respectively. *r*(*z*), the radius as a function of height, was simply determined using similar triangles:

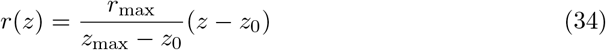

*N*_*s*_(*z*), which describes the linear spine density as a function of height, was assumed a parabola as an approximation of the linear spine densities found in [8] and [40]:

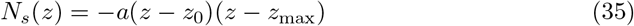

Where *a* is a constant to be determined as follows. Since *N*_*s*_(*z*) is subject to the constraint:

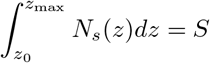

where *S* is the total number of spines on the cone, then:

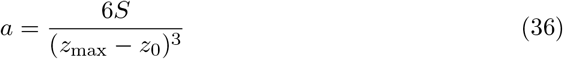

And Eq 35 thus becomes:

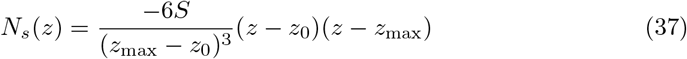

Ultimately, substituting in Eq 34 for *r*(*z*), Eq 37 finally becomes 3:

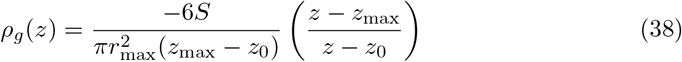

### 0.3 Calculating the overlap dendritic length

From Eq 26, *L*, the total length of dendrite contained in the overlap between an MC and GC, is:

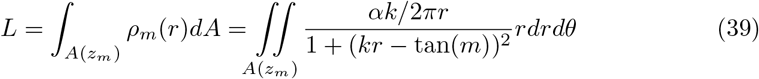

Calculating this integral is dependent on *s*, the distance between the center of the MC field and the GC field, as well as on *r*_*m*_, the radius of the MC, and *r*_*g*_, the radius of the GC at *z*_*m*_, the MC height and thus the height of the intersection. Below we detail how the integral changes as the two fields are drawn closer together.

For all cases, if *s* ≥ *r*_*g*_ + *r*_*m*_, or if *z*_*m*_ is out of range of the GC cone (*i*.*e*. the MC and GC do not overlap), then by default:

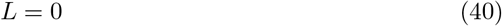

Again for all cases, if 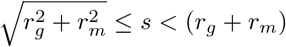 , then

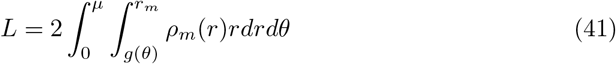

where:

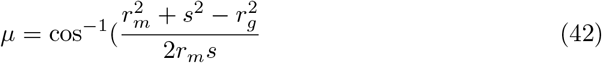

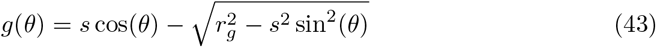

the latter of which is just the equation in polar coordinates for a circle a distance *s* from the origin. Since the problem is symmetric, we multiplied the integral by 2 and integrated from 0 rather than integrate from −*µ* to *µ*. Note that we adopted this strategy for all integrals except the case where the GC field overlapped the center of the MC field, in which case we integrated over all *θ* (see below).

Moving forward, the integrals depended on the relative sizes of *r*_*g*_ and *r*_*m*_, so below we consider the relevant cases separately. Before continuing, we must define two further quantities:

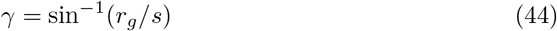

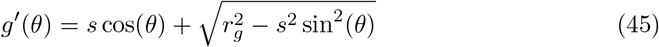

where the latter is again the equation in polar coordinates for a circle a distance *s* from the origin, but integrated in the opposite direction from *g*(*θ*).

**Case 1**: ***r***_***m***_ ***>* 2*r***_***g***_

If 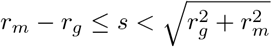

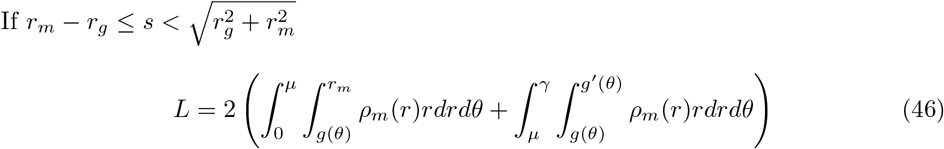

If *r*_*g*_ ≤ *s < r*_*m*_ *− r*_*g*_

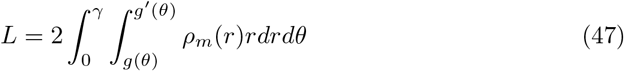

If 0 ≤ *s < r*_*g*_

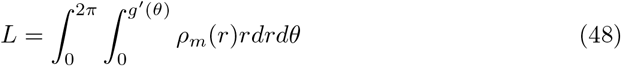

**Case 2**: **2*r***_***g***_ ***> r***_***m***_ ***> r***_***g***_

If 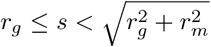

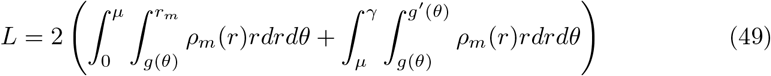

If *r*_*m*_ *− r*_*g*_ ≤ *s < r*_*g*_

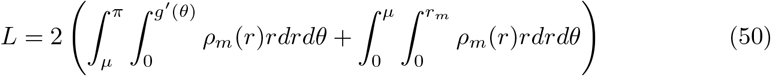

*L* = 2

If 0 ≤ *s < r*_*m*_ *− r*_*g*_

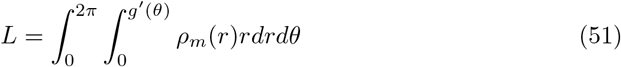

**Case 3**: ***r***_***g***_ ***> r***_***m***_

If 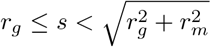

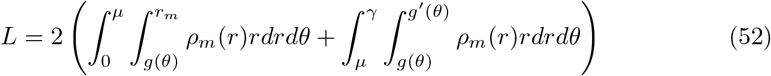

If *r*_*g*_ *− r*_*m*_ ≤ *s < r*_*g*_:

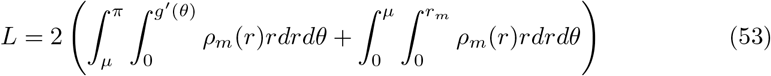

*L* = 2

If 0 ≤ *s < r*_*g*_ *− r*_*m*_:

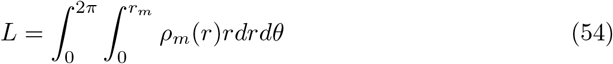

#### Layers of the OB space

We measured the average thickness of the external plexiform layer (EPL), mitral cell layer (MCL), and internal plexiform layer (IPL) from *camera lucida* images in supplementary material from [8] and [9]:

| Layer | Thickness (in $\mu\text{m}$) |
| --- | --- |
| EPL ($\theta_{\text{EPL}}$) | 131 |
| MCL ($\theta_{\text{MCL}}$) | 36 |
| IPL ($\theta_{\text{IPL}}$) | 27 |
| Total ($\theta$) | 194 |

#### Glomerular density and distribution

For the purpose of calculating the area density of the glomerular projections (henceforth referred to as simply ‘glomeruli’) on the EPL, we assumed the EPL to be a flat (with thus no difference in the surface areas of the top and bottom surfaces), 3-dimensional space with a volume of around 1.5 mm^3^ [119]. We also assumed the thickness of the EPL to be uniform (although in general there is considerable variation in thickness over the bulb), and so by dividing this volume by *θ*_EPL_, we arrived at an area of 1.145 × 10^7^ *µ*m^2^. We assumed 1800 glomeruli per olfactory bulb [118, 120], leading to an area density *ρ*_glom_ of 157 glomeruli/mm^2^. Thus, for the network, 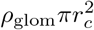 glomeruli were placed uniformly randomly in the x-y plane within a radius of *r*_*c*_.

#### Mitral cell distribution

A number of MCs drawn from Uniform(15, 25) was placed around each glomerulus. For a glomerulus located at (*x*_glom_, *y*_glom_), the location of one of these MCs in the x-y plane was (*x*_glom_ + *r* cos *θ, y*_glom_ + *r* sin *θ*), where *r* (in *µ*m) ∽ Logistic(*µ* = 78.4, *s* = 23.1) for *r* ∈ [0, 300] [4], and *θ* ∽ Uniform(0, 2*π*). The radius of each MC was drawn from Uniform(75, 800).

MCs were either assigned as type I with 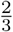 probability or type II with 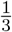 probability [2]. The z-positions for each cell type were drawn from the following distributions:

|  | Type I | Type II |
| --- | --- | --- |
| Z-position | $\theta_{\text{IPL}} + \theta_{\text{MCL}} + \text{Uniform}(0, \frac{1}{2}\theta_{\text{EPL}})$ | $\theta_{\text{IPL}} + \theta_{\text{MCL}} + \text{Uniform}(\frac{2}{5}\theta_{\text{EPL}}, \frac{4}{5}\theta_{\text{EPL}})$ |

#### Granule cell distribution

We specifically modeled deep granule cells, which preferentially interact with mitral cells [8]. GCs were distributed randomly in the xy-plane such that the x and y positions of their bottom vertices were distributed uniformly randomly. The maximum radius, top and bottom z-positions, and xy-distance of the center of the top face from the bottom vertex were drawn from the following distributions (in *µ*m):

|  |  |
| --- | --- |
| Maximum radius ($r_{\text{max}}$) | Normal(83, 28)<br>for $r \in [30, 160]$ |
| Bottom z-position (vertex) | Uniform($0, \theta_{\text{IPL}} + \theta_{\text{MCL}}$) |
| Top z-position (face) | $\theta_{\text{IPL}} + \theta_{\text{MCL}} + \text{Uniform}(\frac{1}{2}\theta_{\text{EPL}}, \theta_{\text{EPL}})$ |
| Vertex to face-center projection distance | Uniform($0, 50$) |

Finally, the number of spines *S* was drawn from a uniform distribution with bounds determined by the volume of the cone. The equation of the bounds was of the form:

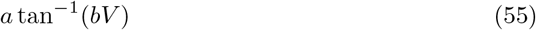

where *V* is the volume. For the lower bound, *a* = 39.31 and *b* = 1.043 ∗ 10^*−*5^, while for the upper bound, *a* = 357.7 and *b* = 2.653 ∗ 10^*−*6^.

### 0.4 Experimental procedures

#### Skewed normal distribution

A skewed normal distribution has a PDF described by the following equation:

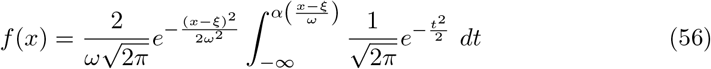

where *ω* is the scale parameter, *α* is the shape parameter, and *ξ* describes the shift of the distribution.

#### Local field potential oscillations

After the LFP signal was calculated, we it through a 6th-order low-pass Butterworth filter with cutoff of 200 Hz and then detrended the signal. We removed the first 200 ms of each of the two periods to roughly isolate the signal’s steady-state for each period. We then used Welch’s power spectral density estimate to calculate the power spectrum, utilizing a window size of 400 ms with 50% overlap between windows.

#### Lateral inhibition

MC pairs (here denoted cells A and B) were selected from then network that had a number of connected GCs within 75 of the average for all MCs and whose z-positions lay within 5 *µ*m of each other. During the first simulation, cell A was fed 700 pA of direct current for 1 s (with 100 ms of unrecorded padding time at the beginning of the simulation to allow the cell to activate from rest), and the consequent firing rate for cell A was measured. During the second simulation, cell A was fed 700 pA of direct current while cell B was fed 750 pA of direct current, and the firing rate for cell A was measured again. This process was repeated for 1,436 different pairs of MCs to cover a wide range of inter-cell distances.

#### Decorrelation

For simulation of a given odor, each MC in each glomerulus received input current of the form:

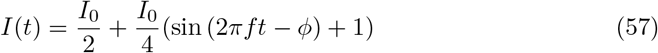

For the MCs belonging to each glomerulus, *I*_0_ was drawn from 𝒩 (*I*_mean_, *I*_mean_*/*5) pA. If the glomerulus was one which was designated to receive odor, *I*_mean_ was drawn from a uniform distribution between 400 and 600 pA, while for all other glomeruli, *I*_mean_ was drawn from a uniform distribution between 0 and 150 pA; the phase *ϕ* was drawn as for the LFP experiments.

Six odors were generated which each targeted 30 glomeruli and which had varying degrees of glomerular overlap (between 5 and 25 shared glomeruli, mean = 18), such that the strength and phase of the input for odor-receiving glomeruli were identical; meanwhile, strength and phase for the different glomeruli and for all other non-odor-receiving glomeruli were different. Odor presentation was simulated for 6 sniffs at 6 Hz (i.e. 1 second of in-simulation time) for each odor individually. Then, the spike time series for each odor was divided into windows of time length *T* , with 50% overlap between windows, and the Pearson correlation was computed between corresponding intervals for each pair of two odors (*n* = 15). The value of *T* was varied to examine how the correlation time course depended on the timescale.

#### Cortical feedback to GCs

In each experiment, an odor was generated which targeted 35 glomeruli. Odor currents were simulated as in the decorrelation experiment, for a total of 2 sniffs (1/3 of a second real-time). For baseline, odor input alone was presented. For the first condition, excitatory feedback was added to a subset GCs in the form of 50 pA of constant current, with the subset of GCs being either 0.1%, 1%, 5%, 10%, 15%, or 20% of all GCs depending on the experiment. For the second condition, feedback was again presented but to a set of GCs non-overlapping with the first set. We examined only the second sniff (since network dynamics appear to stabilize after one sniff) and calculated the change in firing rate that occurred with feedback compared to baseline for each odor-receiving MC for each condition. We computed the Pearson correlation of these changes in firing rates over 10 ms windows, with 50% overlap between successive time windows, between the first and second conditions, and then found the mean correlation. For the third condition, a new arrangement of GCs was generated around the same MCs and the experiment was repeated, with the correlation of the changes in firing rates being computed between the results of the first condition and this new condition.

#### Cortical feedback to MCs

These experiments were conducted similarly to the previous feedback experiments. During the feedback condition, instead of feedback current to the GCs, constant excitatory feedback current was delivered to 0.2 of all MCs, with the strength of the current for each MC being drawn from Normal(200, 20) pA. Data in the figure was compiled from *n* = 5 trials, each with a different odor and feedback pattern.

### 0.5 Simulation

All simulations were done in MATLAB versions R2017, R2018, or R2019 via a forward Euler method with time step = 0.1 ms.

## Supporting Information

**Fig. S1 Distribution of sister MC somata with relation to glomerulus** Distribution of sister MC somata with relation to their parent glomerulus from [4]. This distribution was well fit by a logistic function of the form 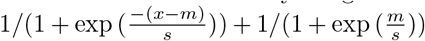, with *m* = 78.4 and *s* = 23.1 (*r*^2^ value = 0.998), and the constant term forcing the function through the origin, since we assume that no MCs are encapsulated by a circle of radius 0.

**Fig. S2 Lateral inhibition and connectivity in the distance-independent network**. (A) When the probability of connectivity is independent of distance, the number of shared GCs (A) and strength of lateral inhibition (B) are also both independent of distance. Distributions of connectivity are Gaussian for both (C) MCs and (D) GCs.

**Fig. S3 Connectivity in the periodic network** Distributions of (A) MC and (B) GC connectivity are right-shifted compared to those of the bounded network.

## Author Contributions

**Conceptualization**: David E. Chen Kersen, Gaia Tavoni, Vijay Balasubramanian

**Formal Analysis**: David E. Chen Kersen

**Funding Acquisition**: Vijay Balasubramanian

**Investigation**: David E. Chen Kersen

**Methodology**: David E. Chen Kersen, Gaia Tavoni

**Project Administration**: Vijay Balasubramanian

**Resources**: David E. Chen Kersen, Vijay Balasubramanian

**Software**: David E. Chen Kersen

**Supervision**: Vijay Balasubramanian

**Validation**: David E. Chen Kersen

**Visualization**: David E. Chen Kersen

**Writing: Original Draft Preparation**: David E. Chen Kersen, Vijay Balasubramanian

**Writing: Review & Editing**: David E. Chen Kersen, Vijay Balasubramanian

## Acknowledgments

DK would like to thank Minghong Ma and Graeme Lowe for helpful discussions, and Yale Cohen, Eugenio Piasini, and Rebecca Li for providing computational resources.

