## Supplementary figures and images for "Connectivity and dynamics in the olfactory bulb"

### Supplemental Figure 1

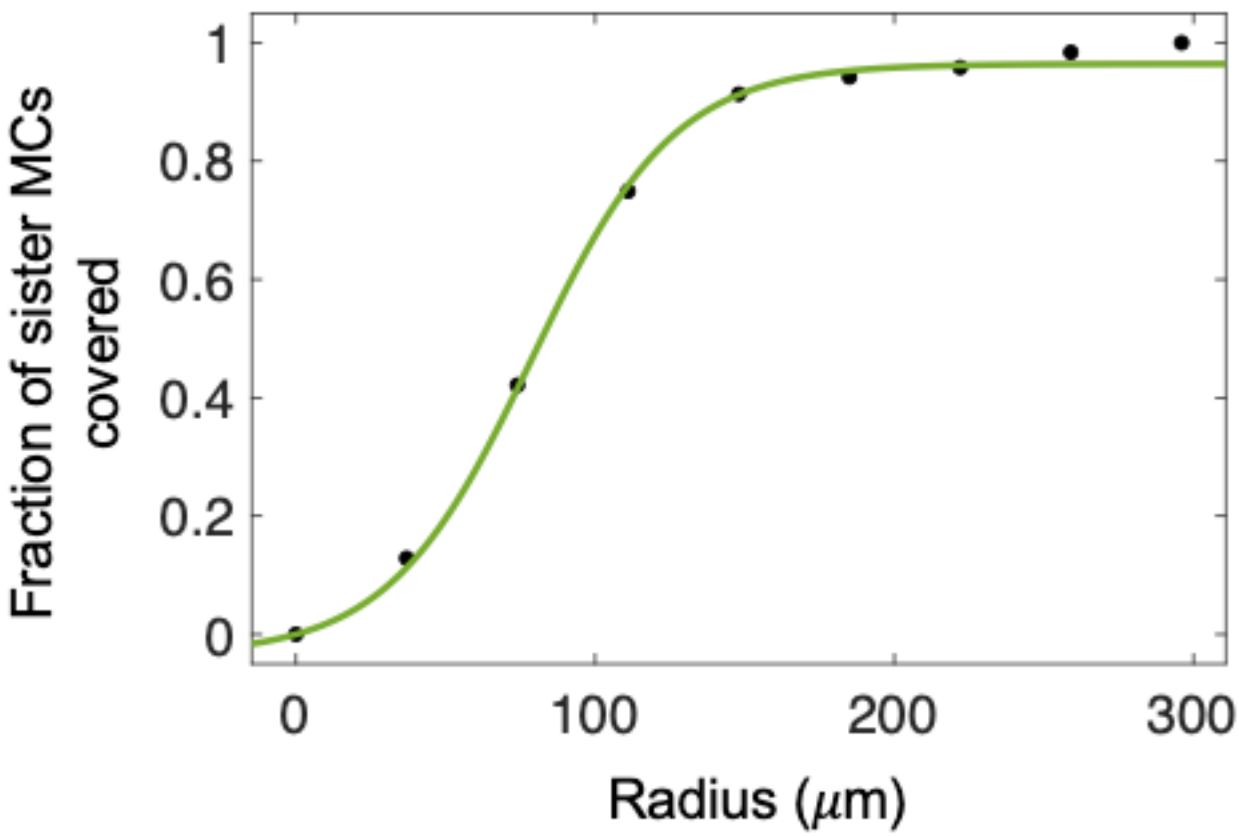

### Supplemental Figure 2

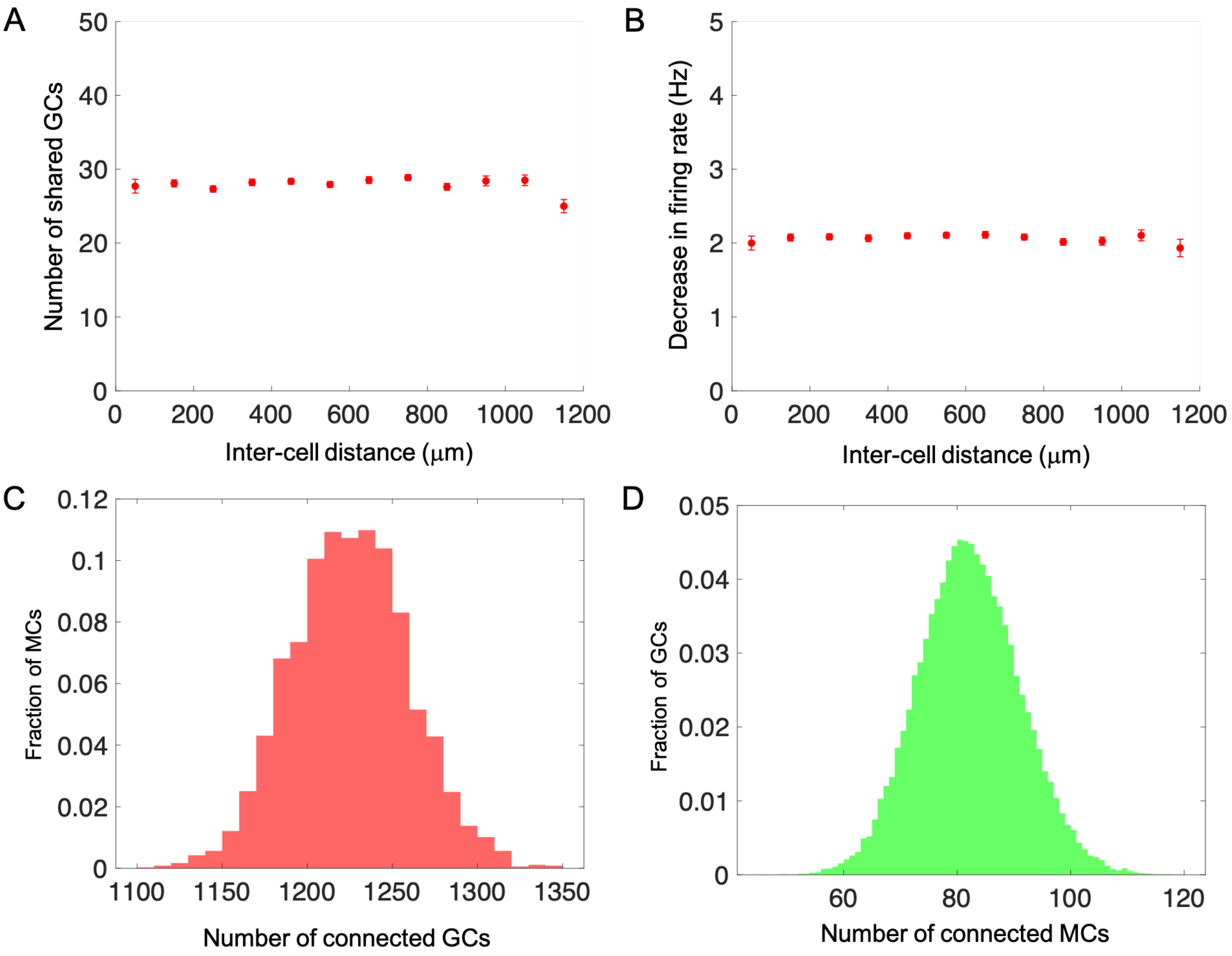

### Supplemental Figure 3

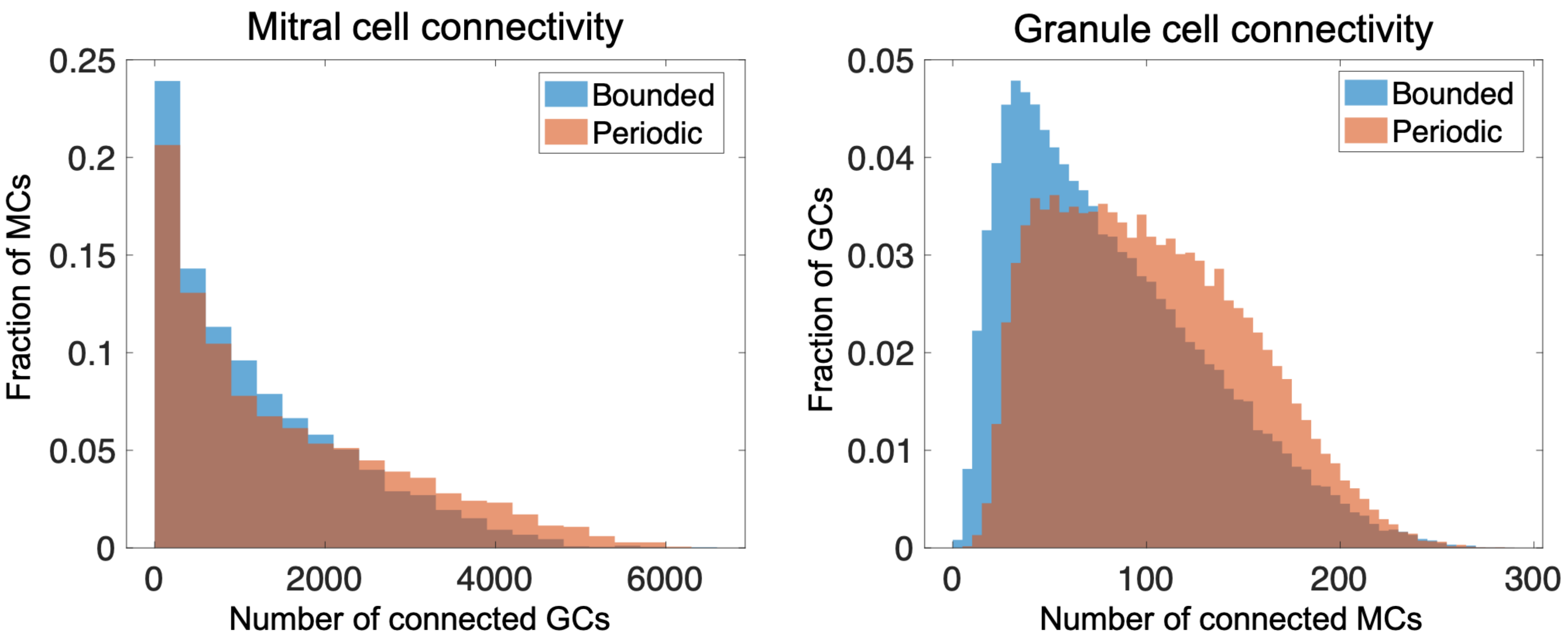
